# Male attention allocation depends on social context

**DOI:** 10.1101/2021.02.27.433194

**Authors:** Shelby D. Burridge, Ingo Schlupp, Amber M. Makowicz

## Abstract

Attention, although limited, is a mechanism used to filter large amounts of information and determine what stimuli are most relevant at a particular moment. In dynamic social environments, multiple individuals may play a pivotal role in any given interaction where a male’s attention may be divided between a rival, a current mate, and/or future potential mates. Here, we investigated impacts of the social environment on attention allocation in male sailfin mollies, *Poecilia latipinna*, which are a part of a sexual-unisexual mating system with the Amazon molly, *Poecilia formosa*. We asked: 1) Does the species of female influence the amount of attention a male allocates to her? And 2) Is a male’s attention towards his mate influenced by different social partners? Males direct more attention toward a stimulus female when she was a conspecific. We also show that males perceive a larger male as a more relevant stimulus to pay attention to than a smaller male, and a conspecific female as a more relevant stimulus compared to a heterospecific female. Our results show differential allocation of attention is dependent upon multiple components of the social environment in which an individual interacts.

**Significance:** This study investigates how attention is allocated in males when presented with social distractions. Assuming that attentional capacity is finite, males may face a tradeoff between different cognitive-demanding stimuli, such as rival males and potential future mates, when mating. Here, we show that male attention allocation in both intra- and intersexual interactions is multifaceted and context dependent. This suggests that individuals within the social environment vary in how meaningful (i.e., able to capture attention) they are to males during mating encounters. Understanding how social partners can cause a shift of attention away from a mating opportunity is essential to understanding the influence of the social context on sexual selection.

## Introduction

Attention is one mechanism that organisms can use to screen the flood of information that is transmitted at any moment in time to determine which stimuli are most relevant at a particular moment (1). However, the ability to filter information and provide selective attention is limited. Attentional capacity is constrained by the size of an organism’s brain and metabolic costs of neural tissue (2). These limits on attention have been shown to have negative effects on the fitness of an individual when attention has to be divided between several tasks. For example, in blue jays, *Cyanocitta cristata*, target detection rates for cryptic food items declined when the birds had to divide attention between a peripheral and central feeding location (3). Detection rates continued to decline when birds were presented with an increasing number of distractor items that divided attention further. Other studies have found a balance between dividing attention towards foraging and predator vigilance. In the presence of a predator, feeding rates often decline as attention is given to predators (4), whereas a division of attention towards high foraging rates may lead to an increased likelihood of succumbing to predation (5). Predators themselves face another trade-off and may be less successful feeders due to constraints on attention. When predators confront large groups of prey items, successful predation rates often decline, as the attention of the predator becomes divided among the individual prey items (6-8).

While there is an increasing body of literature on effects of limited and divided attention on foraging (3), predator avoidance capabilities (4-5), and predation success (6-8), little is known about division of attention in regard to sexual selection. A few studies in humans have shown that sexually unrestrictive men allocated significantly more attention to attractive women and less to unattractive women (9-10). This allocation of attention towards facial attractiveness seems to be dependent upon sociosexuality, relationship status, and perceived self-market value of the choosers (11). Similarly, in animals that live in social groups, several individuals may play a role in any given interaction. When a male is interacting with a particular female, other individuals such as rival males and potential future mates may be present that draw attention away from the current interaction (12-13). In the present study, we investigated how males will divide their attention between a stimulus female and a variable audience to determine how the context of the social environment affects attention allocation, assuming that attentional capacity is finite.

Measuring attention allocation can be challenging in non-primate models where typical methods such as fMRIs (14), event-related potential (11), and eye tracking glasses (15-16) are less feasible options. Most studies, therefore, use other, indirect measurements, particularly the proportion of time allocated to specific tasks (17-18), latency to start a particular task (4,5,17), distance to prey before a predator is detected (19), or the number of correct responses (3,20-21). The time animals spend allocating between two separate tasks (e.g., the tradeoff between predator vigilance and foraging; 18) can be a reliable measure of how they allocate attention and balance their cognitive load. Studying how social distractions influence attention allocation within the context of sexual selection in non-primates can be even more challenging, as it can use the same units of measure (e.g., proportion of time) as in other components of sexual selection (e.g., mate preference). However, even in humans, the proportion of time was used to measure how much attention individuals allocated to attractive opposite-sex faces (9). Thus, we apply this same methodology to study how attention is allocated between the proportion of time males can spend mating with a readily accessible, fertile female and the time he spends interacting with a rival male or a potential, but inaccessible mate. Specifically, we measure the tradeoff between intrasexual competition and mating, where a sexually motivated male is predicted to spend more time physically interacting with a fertile female over visually interacting with a rival male. Similarly, we measure the tradeoff between a readily accessible mate and an inaccessible potential mate, where a sexually motivated male is predicted to spend more time physically interacting with a fertile, accessible female over visually interacting with an inaccessible potential mate.

The sailfin molly, *Poecilia latipinna*, is a species of livebearing freshwater fish that lives in dynamic social groups. Males display sexual behaviors toward preferred females and aggression toward rival males (22). Sailfin mollies provide an interesting system to study attention allocation because in addition to their highly dynamic social environment, they are also host to a sexual parasite, the Amazon molly, *Poecilia formosa*. Amazon mollies are gynogenetic, unisexual hybrids that arose from a natural hybridization event between sailfin mollies and the Atlantic molly, *Poecilia mexicana*; thus, they occur syntopic with their host species (23-24). Male sailfin mollies are capable of distinguishing conspecific females from heterospecific Amazon mollies and show preference towards conspecifics (25-27). The use of the sexual-unisexual mating system of sailfin mollies to address questions regarding effects of the social environment on attention allocation provides another interesting layer of social interaction. By assessing the scenarios in which a sailfin molly male allocates attention towards Amazon mollies, we may be able to further understand how Amazon mollies achieve copulations, which are crucial to them, but mostly costly to the males.

Taking advantage of this sexual-unisexual mating system, we investigated two questions: 1) Does the species of female influence the amount of attention a male allocates to her? And 2) Is a male’s attention towards his mate influenced by different social partners? Specifically, we asked how a focal male divides attention between a stimulus female (either a conspecific or heterospecific female) with whom he can readily interact with and either a rival male which varied in body size or a potential mate (i.e., a conspecific or heterospecific female). To study how intrasexual competition influenced the attention allocation of male sailfin mollies we used eight social conditions: 1) sailfin molly female with smaller-sized male audience; 2) sailfin molly female with larger-sized male audience; 3) sailfin molly female with equally sized male audience; 4) sailfin molly female with no audience; 5) Amazon molly female with smaller-sized male audience; 6) Amazon molly female with larger-sized male audience; 7) Amazon molly female with equally sized male audience; and 8) Amazon molly female with no audience. To assess how potential future mates influence male attention we simulated six social conditions in which a focal male observed: 1) a sailfin molly stimulus female with sailfin molly audience female; 2) a sailfin molly stimulus female with Amazon molly audience female; 3) a sailfin molly female with no audience; 4) an Amazon molly stimulus female with sailfin molly audience female; 5) an Amazon molly stimulus female with Amazon molly audience female; and 6) an Amazon molly stimulus female with no audience.

If attention allocation occurs randomly, we would expect males to split their attention equally between the stimulus female and the audience individual. However, if attention allocation is dependent upon the relative information value of the stimuli, we predicted that males would allocate more attention overall towards the more valuable stimulus female (i.e., the conspecific female), compared to the less valuable stimulus female (i.e., the heterospecific female). In the intrasexual competition social conditions, we predict that the similar-sized males would detract the greatest amount of the focal male’s attention away from the stimulus female (28), there will be little detraction from smaller males (29-30), and males that are larger than the focal male will result in focal males spending less attention towards a stimulus female in order to conceal his preference toward that female and reduce the likelihood that the audience male will copy his mate choice (31-32). For the potential future mate social conditions, we predicted that males would allocate equal amounts of attention when both females are conspecifics, more attention towards the conspecific audience female when paired with a heterospecific stimulus female, and more attention towards the conspecific stimulus female when paired with a heterospecific audience female (25-27).

## Results

### (a) Effects of intrasexual competition on male attention

When analyzing the actual time males spent giving attention to the stimulus female, we found that males spent more time with a stimulus female when she was conspecific and when there was no audience present (Tables 1, 2, *SI Appendix*, Fig. S1, Table S1). This was generally predicted. Males spent the most time interacting with the audience male when paired with a heterospecific compared to a conspecific female. Unsurprisingly, males spent the least amount of time interacting near the Plexiglas container when there was no audience present (Tables 1, 2, *SI Appendix*, Fig. S1, Table S1).

**Table 1:**
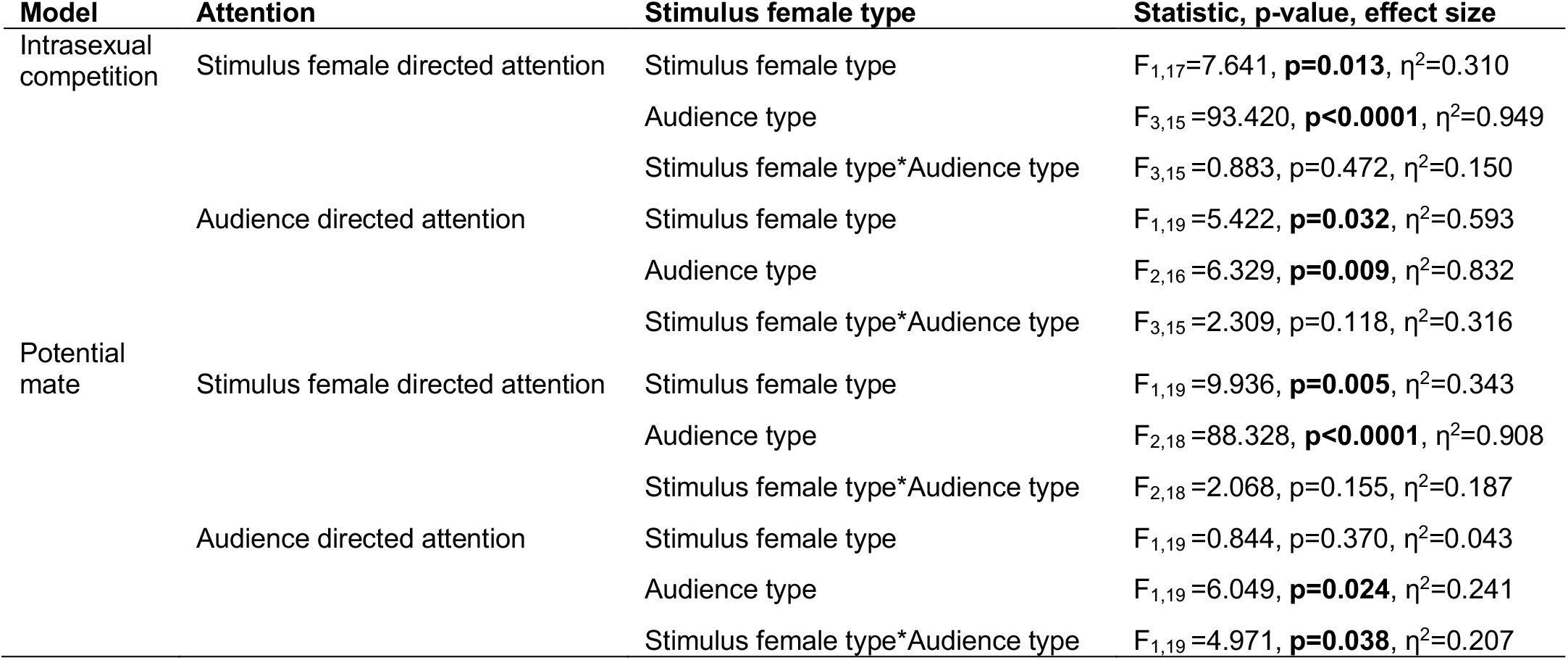
Results from the statistical models on stimulus females and audience directed attention in both the intrasexual competition and potential mate trials.

| Model | Attention | Stimulus female type | Statistic, p-value, effect size |
| --- | --- | --- | --- |
| Intrasexual competition | Stimulus female directed attention | Stimulus female type | $F_{1,17}=7.641$, **p=0.013**, $\eta^2=0.310$ |
| | | Audience type | $F_{3,15}=93.420$, **p<0.0001**, $\eta^2=0.949$ |
| | | Stimulus female type*Audience type | $F_{3,15}=0.883$, p=0.472, $\eta^2=0.150$ |
| | Audience directed attention | Stimulus female type | $F_{1,19}=5.422$, **p=0.032**, $\eta^2=0.593$ |
| | | Audience type | $F_{2,16}=6.329$, **p=0.009**, $\eta^2=0.832$ |
| | | Stimulus female type*Audience type | $F_{3,15}=2.309$, p=0.118, $\eta^2=0.316$ |
| Potential mate | Stimulus female directed attention | Stimulus female type | $F_{1,19}=9.936$, **p=0.005**, $\eta^2=0.343$ |
| | | Audience type | $F_{2,18}=88.328$, **p<0.0001**, $\eta^2=0.908$ |
| | | Stimulus female type*Audience type | $F_{2,18}=2.068$, p=0.155, $\eta^2=0.187$ |
| | Audience directed attention | Stimulus female type | $F_{1,19}=0.844$, p=0.370, $\eta^2=0.043$ |
| | | Audience type | $F_{1,19}=6.049$, **p=0.024**, $\eta^2=0.241$ |
| | | Stimulus female type*Audience type | $F_{1,19}=4.971$, **p=0.038**, $\eta^2=0.207$ |

**Table 2 –.** Descriptive statistics for intrasexual competition effects on male attention according to stimulus female and male audience body size. N = 18 for all groups.

| Stimulus female | Audience male | Time (s) | Mean | Std Dev | Minimum | Maximum |
| --- | --- | --- | --- | --- | --- | --- |
| Amazon molly | Control | Stimulus | 475.07 | 115.80 | 132.44 | 600.00 |
|  |  | Audience | -- | -- | -- | -- |
|  | Smaller male | Stimulus | 326.59 | 87.45 | 137.99 | 439.88 |
|  |  | Audience | 118.38 | 74.30 | 20.04 | 273.76 |
|  | Equal-sized male | Stimulus | 310.77 | 122.89 | 87.52 | 535.12 |
|  |  | Audience | 127.90 | 64.21 | 33.34 | 276.93 |
|  | Larger male | Stimulus | 242.00 | 90.37 | 82.20 | 404.08 |
|  |  | Audience | 195.47 | 105.95 | 59.35 | 438.29 |
| Sailfin molly | Control | Stimulus | 517.95 | 87.82 | 227.45 | 592.34 |
|  |  | Audience | -- | -- | -- | -- |
|  | Smaller male | Stimulus | 376.18 | 98.82 | 158.40 | 502.82 |
|  |  | Audience | 95.44 | 67.92 | 15.72 | 266.92 |
|  | Equal-sized male | Stimulus | 335.76 | 119.76 | 57.50 | 553.31 |
|  |  | Audience | 119.29 | 68.40 | 21.35 | 248.85 |
|  | Larger male | Stimulus | 335.07 | 111.47 | 66.27 | 527.25 |
|  |  | Audience | 137.68 | 77.17 | 25.98 | 327.79 |

When we investigated the proportion of time males spent with either the stimulus female or audience male, we found a significant effect on the direction males allocated their attention (F_1,17_=73.039, p<0.001, η^2^=0.811), with males allocating more attention towards the stimulus female (Fig. 1). Neither the species of stimulus female (F_1,17_=0.900, p=0.356, η^2^=0.050) nor the audience male type (F_2,16_=0.117, p=0.891, η^2^=0.014) caused a significant difference. However, there was a significant interaction between the direction of allocated attention and the stimulus female type (F_1,17_=7.634, p=0.013, η^2^=0.310) and the audience male type (F_2,16_=4.959, p=0.021, η^2^=0.383). Males allocated more attention towards the stimulus female when she was a conspecific compared to a heterospecific (p=0.030) and more attention towards the audience when the stimulus female was a heterospecific compared to a conspecific (p=0.037; electronic supplementary material, table S3). Larger male audiences increased the amount of allocated attention towards the audience male compared to the smaller male audience (p=0.003) or an audience male equal in size (p=0.005). Males directed more attention towards the stimulus female when in front of a smaller male audience compared to a larger male (p=0.020). Finally, there was no significant interaction between the type of stimulus female and the type of audience male (F_2,16_=0.133, p=0.877, η^2^=0.016) or the direction of attention, type of stimulus female, and the type of audience male (F_2,16_=1.536, p=0.245, η^2^=0.161).

**Fig. 1:**
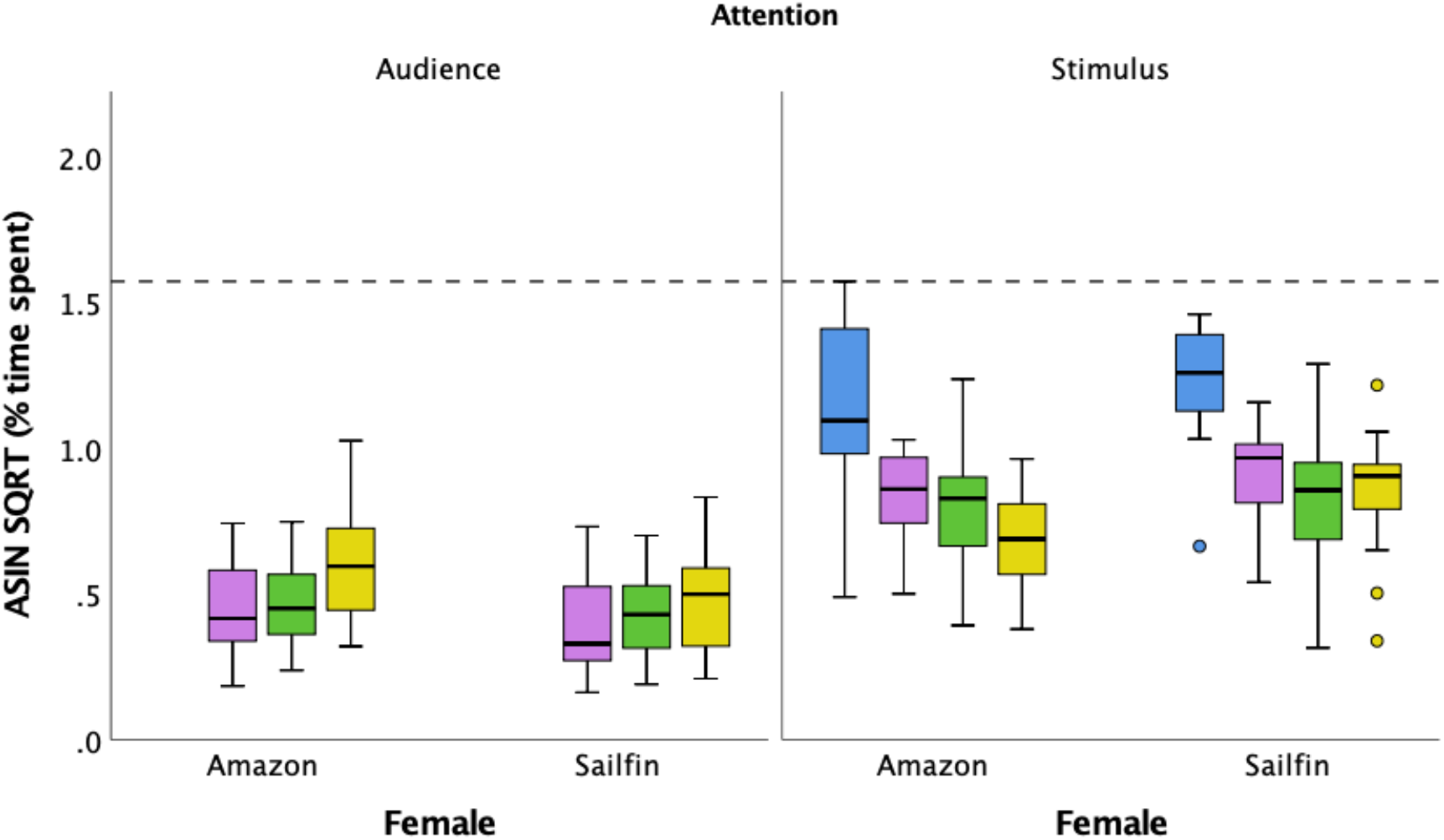
The proportion of time focal males spent with each female when exposed to different male audiences (no audience = blue; smaller male = purple; male equal in size = green; larger male = yellow). Dashed line indicates the maximum amount of time a male could spend with either or both females.

### (b) Effects of a potential mate on male attention

Again, sailfin males spent most their attention towards conspecific stimulus females and spent the most time with the stimulus female when he had no distractions compared to when there was an audience present (Tables 1, 3; *SI Appendix*, Fig. S2, Table S3). Male sailfin mollies directed more attention towards the audience when she was a conspecific female, but there was no effect of the stimulus female type on audience-directed attention. However, there was a significant interaction between the stimulus female type and audience female type, with males spending almost twice as much time with the audience female when that female was a conspecific and the stimulus female was a heterospecific (Tables 1, 3; *SI Appendix*, Fig. S2, Table S3).

**Table 3 –.** Descriptive statistics for female species effects on male attention according to stimulus and audience female species. N=20 for all groups.

| Stimulus female | Audience female | Time (s) | Mean | Std Dev | Minimum | Maximum |
| --- | --- | --- | --- | --- | --- | --- |
| Amazon molly | Control | Stimulus | 504.64 | 71.31 | 339.20 | 600.00 |
|  |  | Audience | -- | -- | -- | -- |
|  | Amazon molly | Stimulus | 276.21 | 107.20 | 43.11 | 425.86 |
|  |  | Audience | 113.02 | 63.99 | 7.76 | 243.79 |
|  | Sailfin molly | Stimulus | 205.50 | 76.16 | 38.74 | 329.76 |
|  |  | Audience | 224.23 | 125.24 | 30.99 | 519.75 |
| Sailfin molly | Control | Stimulus | 531.58 | 102.47 | 252.97 | 660.46 |
|  |  | Audience | -- | -- | -- | -- |
|  | Amazon molly | Stimulus | 315.47 | 131.42 | 110.84 | 527.84 |
|  |  | Audience | 140.43 | 116.41 | 4.75 | 460.13 |
|  | Sailfin molly | Stimulus | 308.02 | 128.83 | 118.59 | 527.84 |
|  |  | Audience | 162.45 | 141.01 | 19.79 | 458.47 |

When we investigated the proportion of time males spent with either the stimulus female or audience female, we found a significant effect of the direction of allocated attention (F_1,19_=23.724, p<0.001, η^2^=0.555) with males spending more attention on the stimulus female (Fig. 2). There was also a significant effect of stimulus female type (F_1,19_=4.922, p=0.039, η^2^=0.206), where males allocated more attention towards conspecific females than heterospecific females. The type of audience female had no significant effect on attention allocation (F_1,19_=1.471, p=0.240, η^2^=0.072). There was a significant interaction between the direction of allocated attention and the type of stimulus female (F_1,17_=4.596, p=0.045, η^2^=0.194), with males spending more time with the stimulus female if she was a conspecific (p=0.019). The direction of allocated attention also had a significant interaction with the type of audience female (F_1,19_=4.574, p=0.046, η^2^=0.194), with males allocating more attention towards the audience female if she was a conspecific female (p=0.021; *SI Appendix*, Table S4). While there was no significant interaction between the type of stimulus female and the type of audience female (F_1,19_=0.609, p=0.445, η^2^=0.031), there was an overall significant interaction between the direction of allocated attention, the type of stimulus female, and the type of audience female (F_1,19_=4.934, p=0.039, η^2^=0.206).

**Fig. 2:**
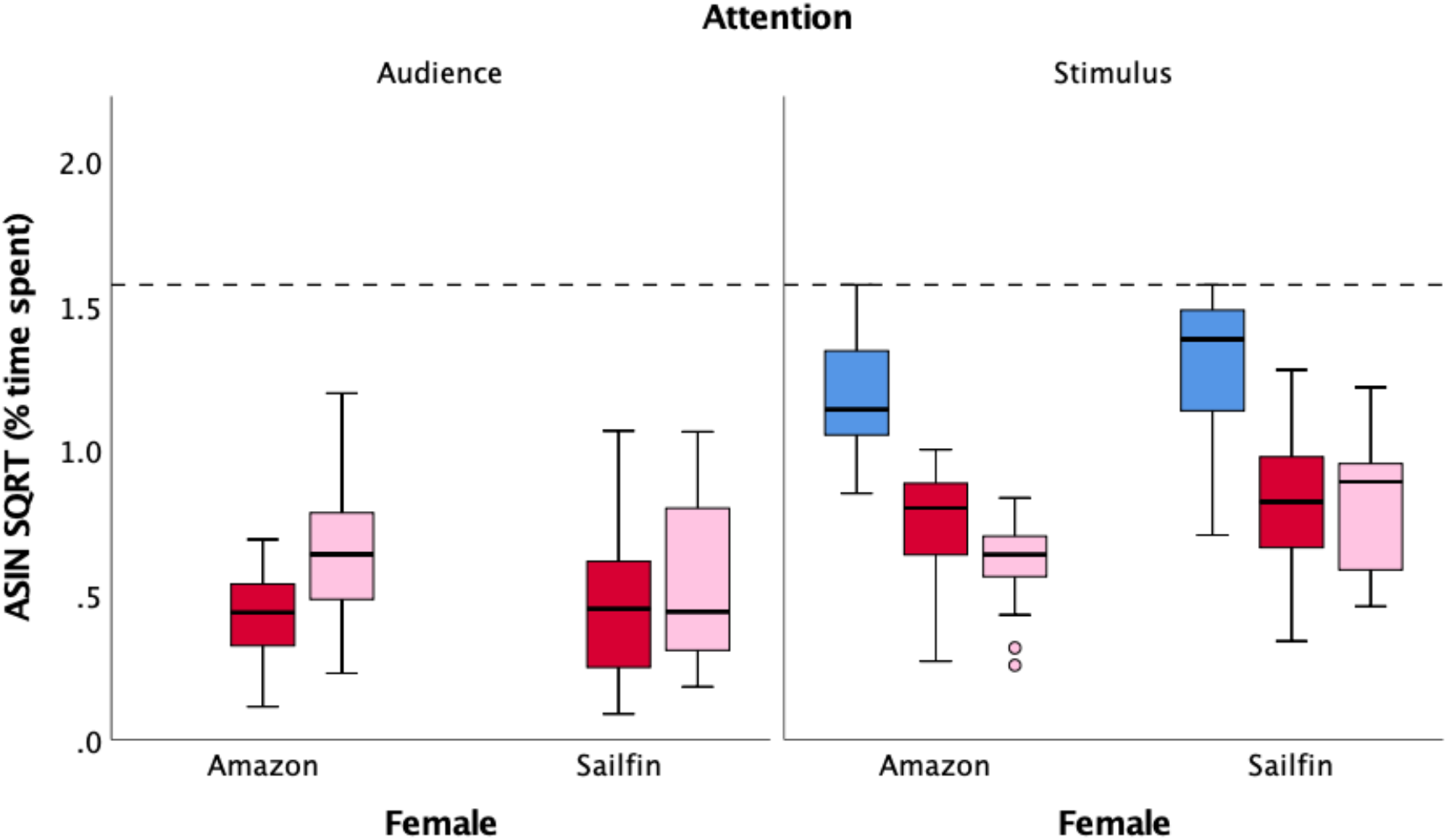
The proportion of attention focal males spent with each female when exposed to different female audiences (no audience = blue; sailfin molly female = pink; Amazon molly = red). Dashed line indicates the maximum amount of time a male could spend with either or both females.

## Discussion

The general presence of an additional social partner, regardless of the sex, size, or species, led to a significant decrease in the attention (measured as time) males allocated to the stimulus female. The general effect of divided attention in both the intrasexual competition and potential mate experiments are unsurprising. When the number of items an individual has to focus on increases, the amount of time and effort an individual can allocate to each is significantly reduced due to neurological constraints (20,33-34). This is especially true here: the amount of time a male allocated towards the stimulus female in the control treatments was significantly higher when compared to any of the audience treatments (Fig. 1, 2). Similar effects of attention divided among multiple individuals have been found in other species. For instance, in Allenby’s gerbils, *Gerbillus andersoni allenbyi*, an individual’s attention may be divided between the presence of a predator and the density load of ectoparasites resulting in a dramatic decrease in time allocated to foraging (18). In the cichlid *Lamprologus ocellatus*, males were shown to divide their attention between predator vigilance and either a conspecific or heterospecific intruding male, and when males were interacting with other conspecific males there was a significant reduction in the distance for males to detect predators (19). Similarly, we show that in the context of sexual selection (i.e., male-male competition and female mate-choice copying), there is also a division of attention between the individuals within the subject male’s immediate social environment regardless of the specific type of individuals. Furthermore, the species of stimulus female is the strongest predictor on how males allocate their attention. In all of the combinations of stimulus female type and audience type, sailfin molly males preferred to interact with conspecific rather than heterospecific females. This result is not unexpected as it has been documented in previous studies that sailfin males prefer conspecific females over heterospecifics (25-26,35). This may also imply that it is more beneficial to the fitness of an individual male to allocate his attention toward a possible mate than to compete with a potential rival or interact with other potential mates.

Contrary to our predictions, males spent significantly more time associating with a larger male than they did with similar-sized males regardless of female stimulus species. Indeed, focal males tended to interact more with the audience when the male was larger than himself compared to either of the other two male size classes. Again, this result is not completely unexpected, given that females within the system prefer larger males over smaller males, and provide more evidence that female choice rather than male competition leads to mating opportunities (28-30). Had the audience and the stimulus male been able to physically interact rather than only being able to interact visually and chemically, these results may have been different. It remains unknown for this study if this switch in attention from the stimulus female to the larger audience male is a way for males to conceal their mate preference or something else altogether (36). Conversely, males appeared to allocate more time toward the stimulus female when the audience male was smaller than the focal male. This result may be due in part to a lower sperm competition risk posed by a smaller male or a male close to the size of the focal male and an increase in male-male competition pressure by a larger male (29-30). However, the result may also imply that males perceive a larger male as a more relevant stimulus than a smaller male, leading to differential division of attention. Finally, association time with the male audience appeared to be influenced by stimulus female type. Males spent more time with the audience male when the stimulus female was a heterospecific than they did when the stimulus female was a conspecific. Since Amazon mollies do not provide a direct benefit to male fitness (23,37), male attention was divided more than when a conspecific female is presented as a stimulus. Nonetheless, these results are evidence that male sailfin mollies are somehow aware of their own body size and can compare his relative size to other males. Furthermore, from this self-reference comparison, males can then alter their attention according to their perceived competition.

There was no overall effect of female audience type on male attention allocation towards the stimulus female beyond the initial decline of adding a second individual. Since the stimulus females are more readily accessible than the audience females (who were enclosed in a chamber), males were able to direct attention towards the stimulus female regardless of audience species in order to gain immediate copulations and increase the possibility that an audience female will copy the mate choice of the stimulus female in future reproductive encounters. While there was no effect of the type of audience female on how much attention males allocated to the stimulus female, there was a significant effect of audience on the amount of attention allocated towards the audience female. Males spent significantly more time interacting with the more valuable conspecific audience compared to a heterospecific audience. This effect was greatest when the stimulus female was a heterospecific Amazon molly.

The results provided by this study show that division of attention is not determined by one factor alone, but rather the components of the social environment in which the organism is interacting. Social context mediates allocation of attention so that an individual gives the greatest amount of attention to the most relevant stimuli at a given moment, whether that is toward a rival male, an accessible female, or a possible future mate. Furthermore, the social partners seem to vary in how meaningful they are to males during mating encounters. Stimuli that have meaning to the individual are better able to capture attention, whereas stimuli that lack meaning fail to (38). Thus, in order to capture attention, upstream cognitive processing must occur to determine the relevance of said stimulus prior to directing attention towards it (39). Our data suggests that the information regarding the body size of a male competitor and the species of a potential mate is being processed by the brain and found to have meaning in the context of current mating encounters (although we did not include a social stimulus that had no meaning to the male). These different social partners also seem to vary in the degree of meaningfulness, as indicated by the differences in attention spent towards each. Understanding what qualities of rival males or potential mates have meaning to males enough to cause a shift of attention away from a mating opportunity is essential to understanding the influence of the social context on sexual selection. In addition, incorporating more social experience and/or social structure into how attention is allocated would provide more information on how the social environment influences social attention and cognition (40). Our study shows that division of attention is multifaceted and context dependent, however, much more work should be done to elucidate a broader range of social implications and the limits of being able to divide attention during sexual and competitive interactions.

## Methods

### (a) Study subjects

Subjects were collected from a drainage basin in Weslaco, Texas (26° 7’13.13”N 97°57’41.08”W) and brought back to the University of Oklahoma, Norman, OK, USA. Collection trips were conducted in May 2015, July 2015, October 2015 and May 2016. Fish were housed in small mixed-sex/mixed-species groups in 37.85-L tanks under 12L/12D light conditions. Fish were allowed to acclimate to laboratory housing for a minimum of 30 days before testing began. After the 30-day acclimation period, measurements of standard length (mm) were taken for all males who were subsequently placed in individual 3.79-L isolation tanks and isolated from females for more than 2 weeks to increase their sexual drive (41). Female fish were then separated by species into 37.85-L tanks to await testing.

### (b) Experimental setup

All trials were conducted in an experimental tank (37.85-L), which had white Plexiglas sheets lining three sides to prevent external distractions. The fourth, long side of the tank remained visible in order to film the behavioral interactions (Nikon D5200 24.1 MP CMOS Digital SLR camera at Full HD (1080p)). A clear, perforated Plexiglas container was placed in the rear center of the tank to house the audience individual (a male or female individual, see below), while two additional perforated Plexiglas containers were placed in the center of the tank to individually house the focal male and stimulus female. After a 10-minute acclimation period, the subject male and stimulus female were released and permitted to interact for 10 minutes. After this period, both the subject male and stimulus female were returned to the Plexiglas containers to await the next trial (see below), or if all trials were complete, both individuals were returned to their former holding tanks. All videos were then analyzed by recording the amount of time (s) a male spent interacting with either the stimulus female or the audience individual.

For this experiment, we assessed how intrasexual competition and potential future mates influenced focal male attention by comparing association times (i.e., the time males spent with the stimulus female compared to the time spent with the audience individual). To assess how intrasexual competition influenced male attention towards the stimulus female, we used audience males who varied in body size relative to the focal male (i.e., a male that was within 2mm of standard length compared to the subject male, a male that was 4mm smaller than the subject male, and a male that was 4mm larger than the subject male). To assess how potential future mates influenced focal male attention, we used an audience female that was either a conspecific (female sailfin molly) or a heterospecific (Amazon molly). Each trial consisted of a subject male paired with either a conspecific or heterospecific stimulus female with an audience individual present. For the intrasexual competition trials, subject males participated in a total of eight social conditions over a two-day period. For the potential mate trials, subject males participated in six social conditions over a two-day period. The two days were split according to stimulus female species, which was chosen at random. On that day, a male would undergo the control for that female species and the three male size trials, which were randomized within that day. All females were size matched to a standard length within ±3mm. We used the same males for both experiments, however, each male was given a one-week rest between experiments.

### (c) Statistical analyses

To assess the time that males allocate to the stimulus female and the audience individual, we evaluated two different data sets for each experiment. In the first data sets, we evaluated the actual time that males allocated to either the stimulus female or the audience individual from the raw data in order to compare the time spent with either individual to the control treatment. Our second data set, we wanted to directly compare the total amount of attention a male allocated towards the stimulus female compared to the audience individual from the total available time. Specifically, we investigated the proportion of time by dividing the raw time spent with each by the total available time (600 s). These values were then arcsin(square root) transformed to fit normality and used in subsequent repeated measures general linear models (GLM).

For the intrasexual competition trials a total of 18 males were tested in each of the 8 treatments, and for the potential mate trials, a total of 20 males were tested in each of the 6 treatments. To assess effects of audience on attention allocation, we ran three repeated measures general linear models (GLM) for the intrasexual competition and potential mate trials. In our first mixed model, male identity was used as a fixed factor to account for variation between individual males while stimulus female type (conspecific or heterospecific) and audience type (intrasexual competition trials: control, smaller than, equal to or larger than focal; potential mate trials: conspecific or heterospecific) were used as fixed factors. The response variable was association time (s) with the stimulus female. In the second model, the random and fixed factors remained the same, while the response variable was association time (s) with the audience individual. The third model tested the proportion of time males spent with either the stimulus female compared to the audience out of the total time available. We used a repeated measures GLM with male identity as the repeated measure, and our fixed factors were attention direction (to stimulus female or audience individual) stimulus female type (conspecific or heterospecific), and audience type. In this model, our response variable was the proportion of time males allocated towards the stimulus female. All models included Bonferroni-corrected post hoc tests to further compare the audience treatments and interactions. All data analysis was conducted using SPSS (IBM v. 24).

## Acknowledgements

We would like to thank Rachel Steele, Elizabeth Hardy, Tana Moore, and Parker Fleming with their assistance on the collection trips and fish maintenance and Texas Parks and Wildlife (permit # SPR-0305-045) for collection permits. This research is in partial fulfillment of MS thesis requirements for SDB.

